# RNA2PS: A sequence-specific coarse-grained RNA model linking structure, thermodynamics and phase separation

**DOI:** 10.64898/2026.04.24.720717

**Authors:** Andrés R. Tejedor, Juan Luengo-Márquez, Javier Oller-Iscar, Jorge Ramirez, Alberto Ocana, Rosana Collepardo-Guevara, Pablo Llombart, Jorge R. Espinosa

## Abstract

RNA plays a central role in the formation and regulation of biomolecular condensates, yet a quantitative understanding of how RNA sequence, structure, and thermodynamics jointly determine phase behaviour, particularly in repeat expansion RNAs, remains incomplete. Here, we introduce RNA2PS, a sequence-specific RNA coarse-grained model for phase-separation that predicts RNA structure from sequence, achieving quantitative agreement for both single-stranded conformations and duplex helical geometry relative to crystallographic PDB structures. RNA2PS represents each nucleotide by two beads that separate the phosphate–ribose backbone from the base. This representation decouples electrostatic interactions from directional base pairing, while explicitly incorporating strand polarity (5′ → 3′) and local sequence context at the trimer level. Canonical and wobble base pairing are modelled through a multi-body potential with sequence-dependent coordination. Importantly, RNA2PS captures sequence-dependent duplex stability at the nearest-neighbour level and reproduces experimental melting temperatures across a diverse set of sequences. RNA2PS shows that phase separation of trinucleotide repeat RNAs is governed by transient inter-strand duplexes that form reversible cross-links. Competition between intra- and intermolecular base pairing regulates the density of labile RNA–RNA interactions, giving rise to strong sequence- and length-dependent differences in condensation that reproduce cellular RNA foci formation. Overall, RNA2PS provides a near-quantitative predictive framework that links sequence-encoded hybridization thermodynamics to mesoscale condensation of pure RNA sequences.

## I. INTRODUCTION

Biomolecular condensates are a fundamental organizing principle of cellular biochemistry^1–3^. These dynamic compartments regulate a wide range of essential biological functions, including ribosome biogenesis in the nucleolus^4,5^, mRNA storage in stress granules^6–8^, and RNA processing and decay in the P-bodies^9,10^. While early studies of biomolecular condensates focused primarily on proteins—particularly those containing intrinsically disordered regions (IDRs)—as the main regulators of condensate biophysics, it is now clear that RNA also modulates condensate structure, composition, and material properties^11–16^.

In RNA–protein condensates, RNA acts as a key modulator of phase behaviour through both specific and non-specific interactions with RNA-binding proteins^17,18^. In particular, it can engage both well-defined RNA-binding domains, such as RNA recognition motifs (RRMs), which provide sequence- and structure-specific binding interfaces, and IDRs enriched in positively charged and polar residues that mediate non-specific favourable electrostatic interactions with the RNA backbone^19,20^. By transforming the intermolecular interactions within RNA–protein condensates, RNA also plays a central role in tuning their material properties^21–24^. For instance, its incorporation has been shown to fluidize condensates of FUS, hnRNPA1 and positively charged peptides^21,22,25,26^, while in other contexts it enhances intermolecular connectivity, promoting more viscoelastic or even gel-like behaviour^23,24,27^.

Beyond its regulatory role in multicomponent protein– RNA assemblies, RNA is capable of undergoing intrinsic phase-separation, without the aid of additional proteins, driven by a combination of electrostatic and sequence-specific interactions^28–31^. The polyanionic backbone of RNA makes electrostatics a central determinant of biomolecular condensate formation via phase separation—herein condensation—with screening by divalent cations such as Mg^2+^ facilitating assembly by reducing phosphate–phosphate repulsion^28,32,33^. At the same time, base-dependent interactions—including *π*– *π* stacking, hydrogen bonding, and cation–*π* contacts— introduce an additional layer of specificity, promoting sequence-dependent cohesion and structural heterogeneity within condensates^34–36^. In particular, RNA repeat expansion sequences provide a well-defined example of this behaviour, where both sequence composition and length govern condensate formation^28^. Trinucleotide and hexanucleotide repeats such as (CAG)_*n*_, (CUG)_*n*_, and (G_4_C_2_)_*n*_ undergo phase-separation above a critical repeat length and concentration^29,30,33^, forming condensates that can evolve from liquid-like droplets to more rigid, gel-like assemblies at high concentrations, a transition directly linked to several neurodegenerative disorders^32,37–40^.

Molecular dynamics (MD) simulations have become essential to unravel the microscopic interactions underlying biomolecular condensation, offering a level of detail that complements experimental observations across multiple length and time scales^41–47^. Atomistic MD simulations with explicit solvent have shown to capture the conformational ensemble of RNA molecules, ionic screening, and specific intermolecular contacts with high fidelity^45,48–51^. Because the high computational cost of atomistic approaches limits their predictive power for biomolecular condensates, coarse-grained models that preserve essential sequence-dependent information are widely used to access the larger system sizes and longer timescales relevant to these systems^52–58^. Within these sequence-dependent coarse-grained models, a simple representation of RNA is to treat it as a generic polyanionic flexible or semi-flexible polymer that primarily captures charge-driven interactions, while largely neglecting sequence specificity, secondary structure, and base-dependent interactions^55,59–61^. The recent CALVADOS-RNA model^55,62^ is a sequence-independent, two-bead-per-nucleotide representation of disordered RNA that separates charged backbone interactions from base-mediated short-range interactions, enabling simulations that capture general aspects of RNA–RNA and protein– RNA interactions while remaining agnostic to RNA sequence and persistent secondary structure.

A notable advance in coarse-grained RNA modelling for biomolecular condensates has been the explicit incorporation of sequence-dependent base-pairing interactions in the single-interaction-site (SIS) model^63,64^. The SIS model represents each nucleotide as a single bead and employs a many-body potential to encode canonical base pairing, enabling simulations of sequence-dependent RNA condensation under effectively charge-neutral conditions. Simulations are typically performed using low-friction Langevin dynamics to accelerate sampling, which is advantageous for exploring the slow kinetics of RNA clustering. Building on this framework, the many-body base-pairing potential was reformulated with analytical forces and implemented in LAMMPS^65^, significantly improving computational efficiency and enabling systematic investigations of sequence-dependent clustering across RNA repeat sequences. While these SIS-based approaches provide a powerful framework for studying sequence-dependent, protein-free RNA condensation, their accelerated dynamical scheme make them not directly compatible with current sequence-dependent protein coarse-grained models for biomolecular condensates, such as the Mpipi, CALVADOS and HPS families^52,57,60,61^, without additional reparameterisation, thereby restricting its applicability to pure RNA condensation ^63,65^. At a resolution intermediate between minimal one- or two-bead models and atomistic descriptions, the iConRNA coarse-grained model^66^, which employs multiple beads per nucleotide and explicit ions, has demonstrated improved accuracy in reproducing RNA conformations, folding, and condensation. However, this increased level of detail comes at a significant computational cost, complicating its application for simulations of multicomponent RNA–protein condensates.

Together, these advancements set the stage to develop RNA models that simultaneously capture sequence-dependent interactions and realistic structural behaviour, while remaining computationally efficient and compatible—in terms of resolution, interaction range, and dynamical timescales—with established coarse-grained protein frameworks for biomolecular condensates^60,67,68^ . In this context, we introduce RNA2PS (RNA to Phase Separation), an RNA coarse-grained model for biomolecular phase separation designed to operate at a comparable level of coarse graining, with interaction potentials amenable to integration with protein models such as Mpipi, CALVADOS, and HPS^52,54,55,57,60,61,69^. RNA2PS is expected to enable applications to multi-component RNA–protein condensates, although this remains to be systematically assessed.

RNA2PS (RNA-to-Phase-Separation) is a coarse-grained model designed to capture the essential structural and thermodynamic properties of RNA within a minimal two-site-per-nucleotide representation. The model considers implicit solvent and incorporates explicit backbone connectivity together with effective base– base interactions that account for *π*–*π* stacking, hydrogen bonding, and directional base pairing, enabling the formation of both single-stranded and double-helical conformations in a fully dynamic manner. Inspired by modern sequence-dependent coarse-grained models of DNA^70–72^, RNA2PS incorporates local sequence specificity by modulating base-pairing interactions according to the nearest-neighbour context up to the trimer level. This local parameterisation captures the influence of adjacent bases on stacking, steric constraints, and binding competition, thereby encoding cooperative effects that are essential for reproducing both local structural features and emergent collective phase behaviour. The model is quantitatively calibrated against experimental data at multiple levels: it reproduces RNA structural features from crystallographic PDB structures and captures sequence-dependent melting temperatures across a diverse set of sequences, including mismatches. In addition, RNA2PS accurately reproduces RNA condensation propensity across different sequences, including repeat expansion systems, capturing experimental trends in condensation and phase behaviour. In particular, the model predicts critical temperatures for RNA phase-separation within biologically relevant ranges and with significantly improved accuracy compared to previous approaches for RNA repeat condensates. By bridging structural fidelity, thermodynamic accuracy, and computational efficiency, RNA2PS provides a unified framework to investigate RNA-driven phase separation and establishes a foundation for future studies of RNA-containing biomolecular condensates.

## II. RESULTS

### A. RNA2PS: A two-site coarse-grained model for RNA condensation

We present RNA2PS, a coarse-grained MD model of RNA designed to capture RNA self-assembly and condensation. RNA2PS employs a two-bead-per-nucleotide representation, in which one bead is placed at the geometric centre of the phosphate–ribose group (hereafter phosphate bead, P), while the second is positioned 4 Å along the vector connecting P to the centre of the nitrogenous base (hereafter base bead, B). In contrast to previous models^63,65^, this representation explicitly decouples electrostatic interactions—localized on the phosphate beads—from the interaction sites responsible for base pairing and cross-stacking, which are carried by the base beads. This separation introduces additional structural flexibility, enabling an accurate description of RNA secondary structures as well as local conformational features of the RNA backbone in single-stranded chains.

In RNA2PS, the intramolecular primary structure is defined through bonded interactions between consecutive phosphate beads (P–P), between phosphate and base beads within the same nucleotide (P–B), and between bases of adjacent nucleotides along the backbone (B– B), effectively encoding the phosphodiester connectivity of the chain (see Section IV and Fig. 1A). Backbone flexibility is primarily controlled by a harmonic angular potential between P beads, parametrised to reproduce the characteristic conformational ensemble of single-stranded RNAs. Within this framework, RNA2PS supports the formation of both intra- and inter-chain base pairs, including canonical (A–U and C–G) and wobble (G–U) interactions, modelled via a many-body potential^63^ (Fig. 1B, top). Base-pair formation is further coupled to angular and dihedral constraints that promote double-helical geometry upon binding, in contrast to the more flexible nature of single-stranded RNAs (Fig. 1B, top).

**FIG. 1.**
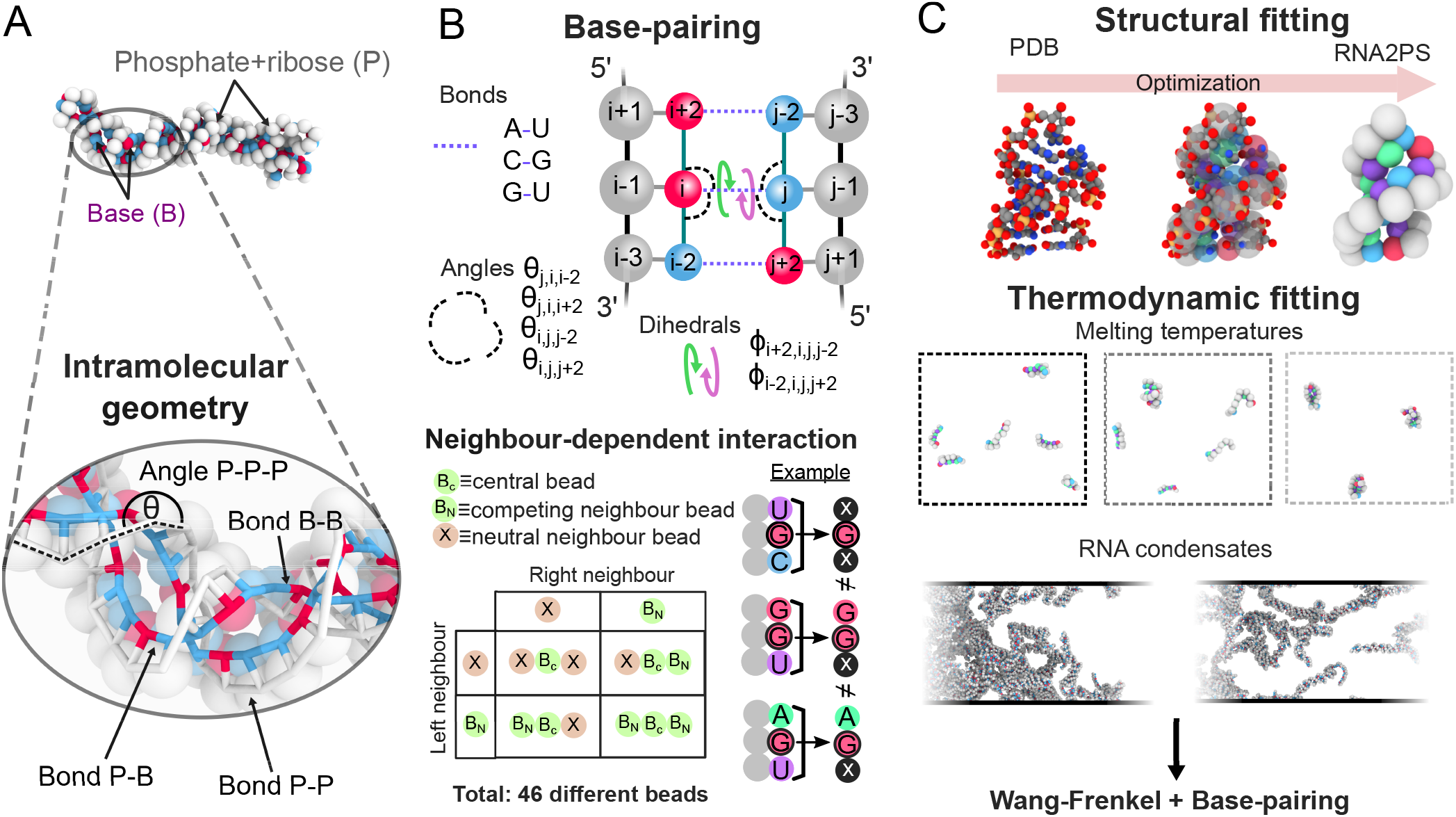
RNA2PS model to describe RNA phase behaviour with two-bead-per-nucleotide resolution. **A**. (Top) Sketch of the RNA2PS model intramolecular interaction sites to describe the RNA structure. The phosphate+ribose bead (P) is represented in gray and a generic base bead (B) is depicted in blue and red. (Bottom) Zoom on the intramolecular interactions sustaining the geometry of double-stranded RNAs, including the bonds and the angle of the backbone. **B**. (Top) Breakdown of the multi-body potential inducing RNA base-pairing. This includes the different bonds, angles, and dihedrals. (Bottom) Schematic table showing the implemented interaction depending on the nature of the nearest neighbours. We present the different four cases considering a central bead of a triad (B_C_) with neutral neighbour (X) and competing neighbour (B_N_) combinations. We provide an example with G with no competing neighbours (C, U), with one G neighbour (binding to C and U) and with one A neighbour (binding to U). In Table S1 we provide the full list of interacting sites. **C**. (Top) We use PDB-based experimentally resolved crystal structures (*e*.*g*. 4JAB) to fit the geometric parameters of both intramolecular and base-pairing interactions. (Bottom) We fit experimental melting temperatures (*e*.*g*. GUAUAUAC) and condensation propensity of RNA repeats (*e*.*g*. CAG_40_) to obtain the interaction strength of the base-pairing multi-body potential and van der Waals (Wang-Frenkel potential) interactions, respectively.

The effective interaction strength of each base pair is determined not only by the identities of the interacting nucleotides, but also by their local sequence context up to the trimer level (see Fig. 1B, bottom). In particular, the interaction energy is modulated by the identity of the nearest neighbours flanking each base, distinguishing between neutral neighbours that do not compete for binding and competing neighbours that introduce alternative pairing configurations (please see Table S1 for the full list of nucleotide types and their interaction parameters). This context dependence encodes an additional layer of effective cooperativity and competition, whereby neighbouring bases can either stabilise or frustrate pairing, thereby promoting collective binding along the RNA strand. Importantly, it also enables RNA2PS to distinguish strand polarity 5^*′*^–3^*′*^, favouring antiparallel base-pairing geometries in a physically consistent manner. This strategy is conceptually inspired by state-of-the-art sequence-dependent coarse-grained DNA models such as the CGeNArate^71^ and MADna^70^ models, where local sequence context is incorporated to reproduce structural and mechanical properties with high fidelity. In RNA2PS, this principle is extended to RNA condensation, enabling the model to capture how subtle sequence variations propagate to large-scale conformational and thermodynamic properties while retaining computational efficiency.

In addition to Watson–Crick base pairing, RNA strands can interact through non-canonical pairing^73^ and base–base stacking interactions^74^. In RNA2PS, stacking interactions between non-adjacent bases are captured using a Wang–Frenkel potential^75^, which depends on the distance between base beads and their biochemical identity. Importantly, this potential also encodes an intrinsic directionality bias that favours correct 5^*′*^–3^*′*^ alignment between interacting strands, thereby stabilising antiparallel configurations consistent with canonical RNA duplex geometry. This interaction plays a dual role: it drives the condensation of RNA sequences that do not form explicit canonical base pairs, and modulates base pairing by both competing with canonical interactions— through the stabilisation of alternative contacts—and facilitating base-pair formation by promoting favourable spatial proximity, orientation, and registry between interacting bases.

The parametrisation of RNA2PS follows a hierarchical strategy. First, structural parameters are optimised to reproduce fundamental topological features derived from experimentally resolved RNA crystal structures (Fig. 1C), while higher-order structural properties, such as specific dihedral distributions, emerge without explicit fitting (Section II B). Second, interaction parameters governing base-pairing and stacking (Wang–Frenkel) are tuned to reproduce experimentally measured RNA duplex melting temperatures (Section II C) and the association propensity of RNA repeat sequences that do not form canonical base pairs (Section II D). As discussed in Section II C, this parametrisation enables the model to quantitatively reproduce experimental trends in phase separation of base pairing RNA repeat sequences, as well as the dependence of condensate stability on sequence repeat length. In the present parametrisation and applications (Section II D), electrostatic interactions, modelled via a Yukawa potential, are neglected to mimic high-salt conditions (e.g., in the presence of Mg^2+^) under which pure RNA phase-separation is observed^28^. Nevertheless, the modular design of RNA2PS—combining electrostatic and sequence-specific dispersive interactions—makes it readily extensible to the study of RNA–protein mixed condensates under physiological conditions.

### B. Structural parametrisation and validation of RNA2PS for double-stranded helical RNA

The conformational properties of RNA play a central role in shaping its interaction landscape and its propensity to form condensates, with features such as chain flexibility, persistence length, and secondary structure directly influencing intra- vs. intermolecular interactions^48,51,76^. Although the primary goal of RNA2PS is not high-resolution structural prediction, as in models such as oxRNA^74,77^, it remains essential to capture a physically realistic spectrum of RNA conformations, ranging from semiflexible single-stranded sequences to double-stranded helical conformations. In this context, recent structure-prediction approaches such as FARFAR2^78^, trRosettaRNA^79^, and RoseTTAFoldNA^80^ high-light the central role of base-pairing and stacking interactions in determining RNA structure, as well as their contribution to intermolecular association and condensation.

The RNA2PS model parameters are determined through a structure-based optimization procedure aimed at reproducing experimentally resolved RNA conformations from PDB structures (see Table S1 in the Supplementary Material for the codes of all the PDBs used). The workflow consists of two main steps: a coarse-grained mapping followed by parameter fitting (Fig. 2A). In the mapping stage, each nucleotide is represented by a minimal set of interaction sites, with the P–B bond length fixed at 4 Å to ensure structural stability of double-stranded RNA. This choice yields an average P–P distance of 15 Å along the helix, preserves consistent P–B– B–P geometries, and prevents unphysical strand crossing. In the fitting stage, a diverse set of RNA structures is considered, including canonical and wobble base pairs (*e*.*g*. 4JAB, 2L2J codes), as well as structural motifs such as hairpins and bulges (e.g. 1A4D, 2A43, 1DQF).

**FIG. 2.**
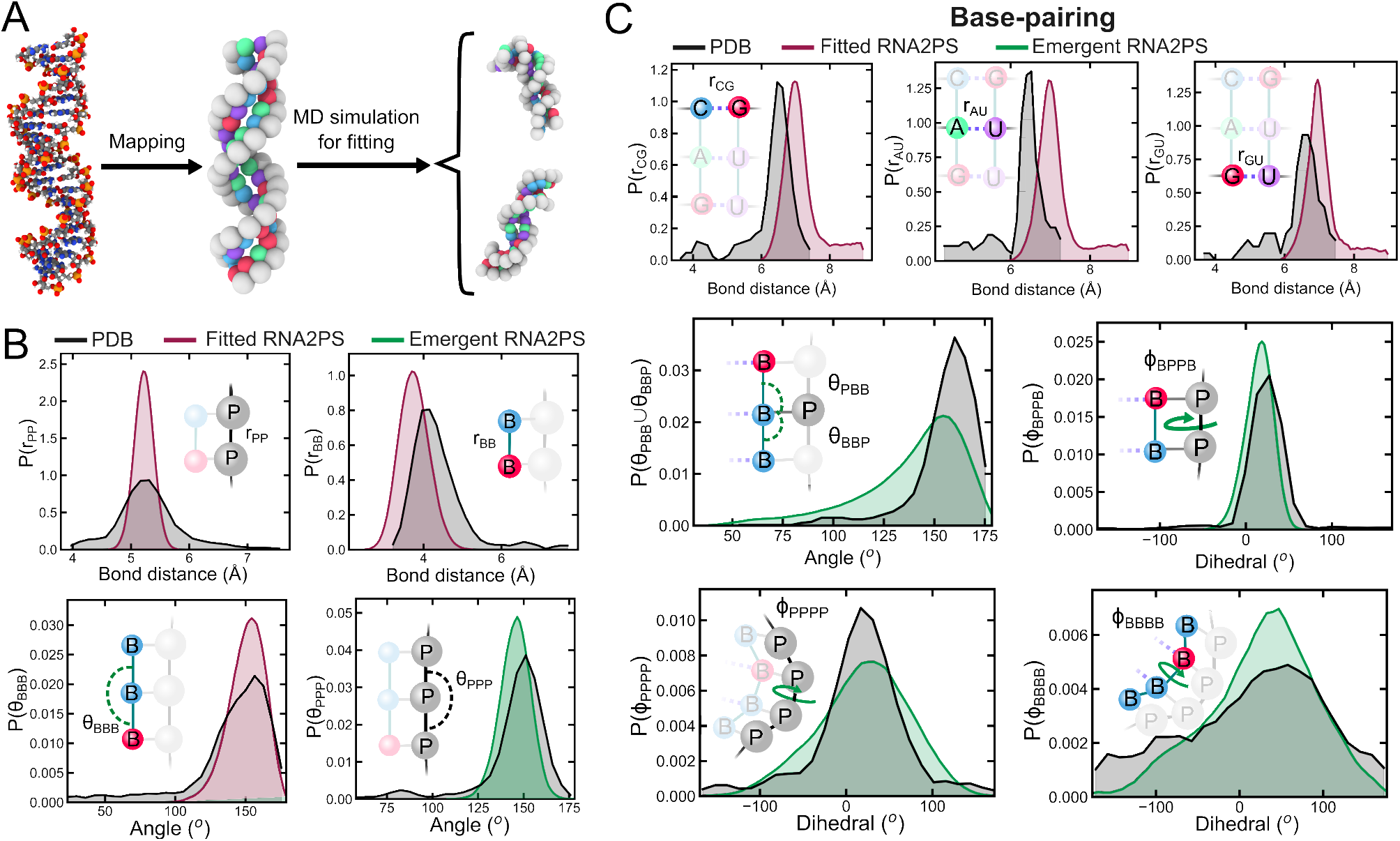
Structural fitting of the model to crystal structures to recapitulate single-vs. double-stranded RNA conformations. **A**. Schematic representation of the structural fitting starting from the all-atom PDB, mapping to the 2-bead-per-nucleotides model and performing molecular dynamics simulations to obtain the statistics about molecular conformation. **B**. Normalized distributions of the intramolecular structure of RNA2PS including the explicitly parametrised and fitted bonds (P-P and B-B) and angles (B-B-B) in red, and the emergent geometric feature of P-P-P angle in green, compared to their analogue calculated from the PDB structures after mapping in black. **C**. Normalized distributions of local conformational parameters related to the base-pairing for the RNA2PS model. Red histograms represent the structural parameters explicitly included and fitted in RNA2PS (*i*.*e*. CG, AU and GU bonds), green distributions are obtained for emergent geometric properties from base-pairing in the RNA2PS model (*i. e*. BBP/PBB angles; and BPPB, PPPP, and BBBB dihedrals), and black histograms show the statistics for the homologue parameters computed from the crystal PDBs.

These structures are simulated and systematically compared to their corresponding crystallographically conformations, allowing the model interaction parameters to be optimized to reproduce structural features of both single- and double-stranded RNAs.

The resulting intramolecular structure shows that the model accurately reproduces key geometric features of RNA (Fig. 2B). In particular, the *r*_PP_ and *r*_BB_ distance distributions, together with the *θ*_BBB_ angle, correspond to observables that are directly fitted and explicitly defined in the model (see Section IV). In contrast, the *θ*_PPP_ angle (expected to be 152°) is not explicitly parametrised and instead emerges naturally from the model topology, while still being well reproduced. These structural descriptors are computed over nucleotides in both single-stranded and base-paired regions, ensuring a consistent description across different conformational states. Notably, the *r*_BB_ distribution obtained from simulations exhibits a slight shift toward more compact conformations compared to the experimental crystallographically resolved structures, consistent with the model design aimed at promoting double-helical formation. Finally, since the P–B bond length is fixed at 4 Å during the mapping procedure, its distribution is reported from simulations in the SM (Fig. S1).

Regarding base-pairing interactions, we analyse the bond length distributions for canonical (CG and AU) and non-canonical wobble (GU) pairs (Fig. 2C). The resulting distributions exhibit slightly larger equilibrium distances compared to atomistic references, reflecting the dynamic nature of pairing imposed by the many-body potential, which operates beyond the excluded volume defined by van der Waals interactions (modelled here via the WF potential). To ensure a consistent characterisation, statistics are computed from MD simulations only when bases are paired—requiring at least one base to be paired for angle distributions and both bases for dihedral angles. Importantly, several structural features emerge naturally from the model. In particular, the *θ*_PBB_ (or equivalently *θ*_BBP_) angles and the *ϕ*_BPPB_ dihedral are accurately reproduced, the latter being essential for capturing the correct helical twist of double-stranded RNAs^51^. Similarly, the backbone dihedral *ϕ*_PPPP_ and the base-stacking dihedral *ϕ*_BBBB_ show good agreement with reference structures, despite their intrinsic flexibility. Notably, RNA2PS explicitly parametrises angular and dihedral terms only for paired bases (see Fig. 1B and Section IV), while the remaining structural correlations emerge implicitly, highlighting the model’s ability to capture key geometrical RNA features with a minimal set of explicit constraints.

### C. RNA2PS captures sequence-dependent RNA duplex stability via melting temperatures

Accurately predicting the thermodynamic stability of ds-RNA is essential for describing its conformational behaviour and its propensity to form pure homotypic condensates^81^. In RNA, this stability arises from a subtle interplay between base-pairing and stacking interactions, which together govern the formation and disruption of double-helical structures^82,83^. To calibrate these contributions, RNA2PS is parametrised against experimental melting temperatures (*T*_m_), which provide a direct measure of RNA duplex stability. Specifically, we consider a dataset of 33 ds-RNA sequences of varying lengths, including five sequences containing mismatches^30,84^.

Fig. 3A shows the time evolution of the fraction of RNA single strands for the representative sequence GUAUAUAC at six different temperatures. At sufficiently high temperatures (dark red curve; 355 K), strands remain predominantly unpaired, whereas at low temperatures (dark blue curve; 280 K) the system undergoes continuous association and dissociation before reaching a well-defined steady state, characterised by an equilibrium between paired and unpaired populations. In this regime, the melting temperature *T*_m_ is obtained from the steady-state ensemble as the temperature at which 50% of the molecules are in the single-stranded state (see snapshot in Fig. 3A). The model parameters governing base pairing are then optimized to reproduce this equilibrium behaviour across a diverse set of sequences, ensuring a consistent description of thermodynamic stability across different RNA lengths and compositions^84,85^. In Fig. 3B, we report the steady-state fraction of single strands as a function of temperature, showing the expected sigmoidal transition between bound and unbound regimes. For this sequence, the RNA2PS model shows an excellent agreement with experimental melting temperature^86^ (blue curve). In contrast, comparison with the SIS model (red curve)^**?**^ reveals a systematic underestimation of the melting temperature by up to ∼ 100 K. This deviation reflects the more permissive interaction landscape of the SIS framework, which, while facilitating structural rearrangements relevant to RNA condensation, reduces its ability to accurately balance association and dissociation equilibria in short sequence duplexes.

**FIG. 3.**
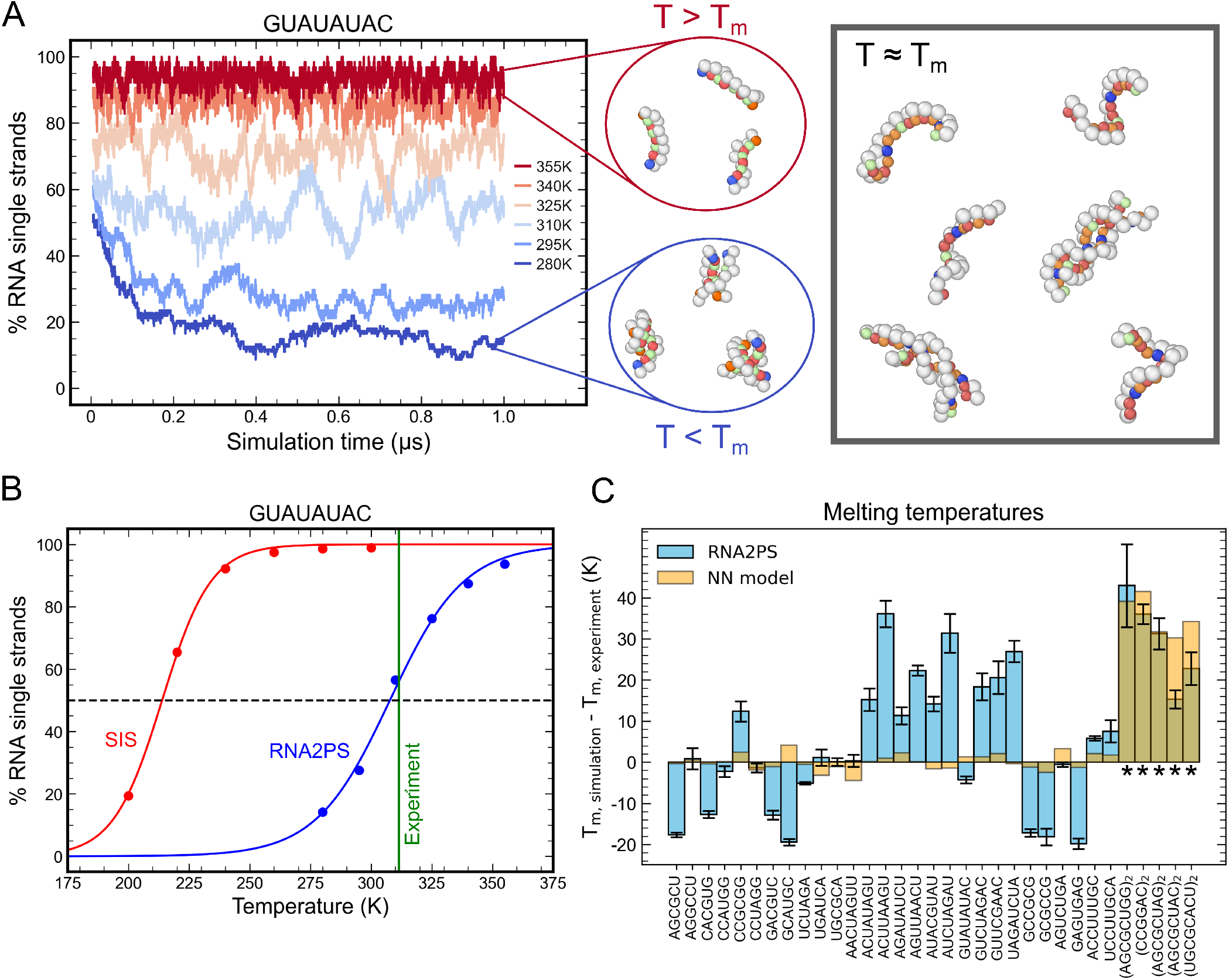
RNA2PS reproduces sequence effects in melting temperatures (*T*_*m*_). **A**. Time evolution of the percentage of RNA single strands for the sequence GUAUAUAC at different temperatures in the range 280–355 K. Representative configurations of the RNA sequences below and above the melting temperature *T*_m_ are shown as insets. We also provide an illustrative snapshot of our system at *T* = *T*_*m*_. **B**. Steady-state fraction of RNA single strands as a function of temperature for GUAUAUAC. RNA2PS results are shown in blue, while SIS model predictions are shown in red. The vertical green line indicates the experimental melting temperature, and the horizontal dashed line marks the 50% unpaired threshold used to define *T*_m_. **C**. Deviations between simulated and experimental melting temperatures for 33 RNA sequences perfectly paired^85^ and with mismatches^84^ (marked with asterisks). RNA2PS results are shown in blue, while the nearest-neighbour (NN) model^84,85^ is shown in yellow.

The bar plot in Fig. 3C summarises the performance of RNA2PS across the full set of 33 RNA sequences by reporting the deviation between simulated and experimental melting temperatures (*T*_m,sim_ − *T*_m,exp_). Overall, the model reproduces both the magnitude and sequence dependence of RNA duplex stability, with an average standard deviation of ∼ 17.5 K across all sequences, indicating robust transferability across different lengths and compositions. While most deviations are centred around zero, small systematic differences are observed for a subset of mismatch-containing sequences, where local destabilization effects are more pronounced. For comparison, the standard nearest-neighbour (NN) model^84,85^, explicitly parametrised for short-duplex thermodynamics^87,88^, achieves slightly lower deviations for short, well-characterised sequences (orange bars), but its performance becomes less consistent outside its optimal parameterisation regime (e.g., the sequences with mismatches marked with asterisks in Fig. 3C). Importantly, those cases are better captured by RNA2PS considering that those sequences were not included in the training of the NN model^84^.

### D. Sequence-dependent condensation in RNA repeat expansions

A central objective of the RNA2PS model is to capture RNA condensation behaviour while preserving a consistent description of RNA thermodynamics and structural accuracy across both single- and double-stranded regimes. In particular, trinucleotide repeat expansions provide a stringent benchmark, as they are known to self-assemble into condensates and gel-like states through multivalent RNA–RNA interactions^28,30^. In RNA2PS, this behaviour emerges from the same interaction framework that governs duplex formation. Base-pairing and stacking interactions, previously calibrated to reproduce melting temperatures across diverse sequences (Fig. 3), act in concert with sequence-dependent cross-interactions encoded through a Wang–Frenkel potential^75^ (see Methods). The latter depends only on base identity and controls the balance between effective hydrophobicity, as well as attraction vs. repulsion in non-base-pair-forming repeats. This unified parametrisation enables the RNA2PS model to reproduce experimentally observed phase-separation trends in RNA repeats (*n* = 40)^30^ without introducing additional fitting parameters for condensation (see Fig. 5C).

To quantify RNA condensation, we perform direct coexistence (DC) simulations of solutions of RNA molecules^89,90^, which enable the explicit observation of coexisting condensed and dilute phases. Below the critical temperature, RNA molecules spontaneously form a dense phase stabilized by dynamic intermolecular base pairing, coexisting with a dilute phase of dispersed strands (see Fig. 4A). This behaviour is characterised through density profiles, from which we extract the equilibrium densities of both phases at different temperatures to construct the temperature–density phase diagram (Fig. 4B). The critical temperature *T*_c_ is obtained by fitting the coexistence curve to the universal scaling law of density differences and the law of rectilinear diameters^91^ (see Section IV). For representative sequences such as CCG_40_ and CGU_40_, RNA2PS predicts critical temperatures close to ∼ 300 K (Fig. 4B), in good qualitative agreement with experimental trends^30,92^. In contrast, simulations with the SIS model yield significantly higher critical temperatures (Fig. 4C), indicating an over-stabilized condensed phase. This discrepancy likely arises from the highly cooperative nature of base pairing in the SIS model, where single-bead nucleotides can simultaneously engage in multiple intermolecular interactions across different strands and orientations, effectively artificially enhancing multivalency and leading to excessive stickiness. In the RNA2PS model, this effect is mitigated by the two-bead representation, which geometrically constrains base pairing and disfavors the formation of multiple simultaneous bonds unless they are coherently aligned along the same strands. As a result, in RNA2PS the balance between inter- and intramolecular interactions is more realistically captured, yielding condensate stabilities within biologically relevant conditions.

**FIG. 4.**
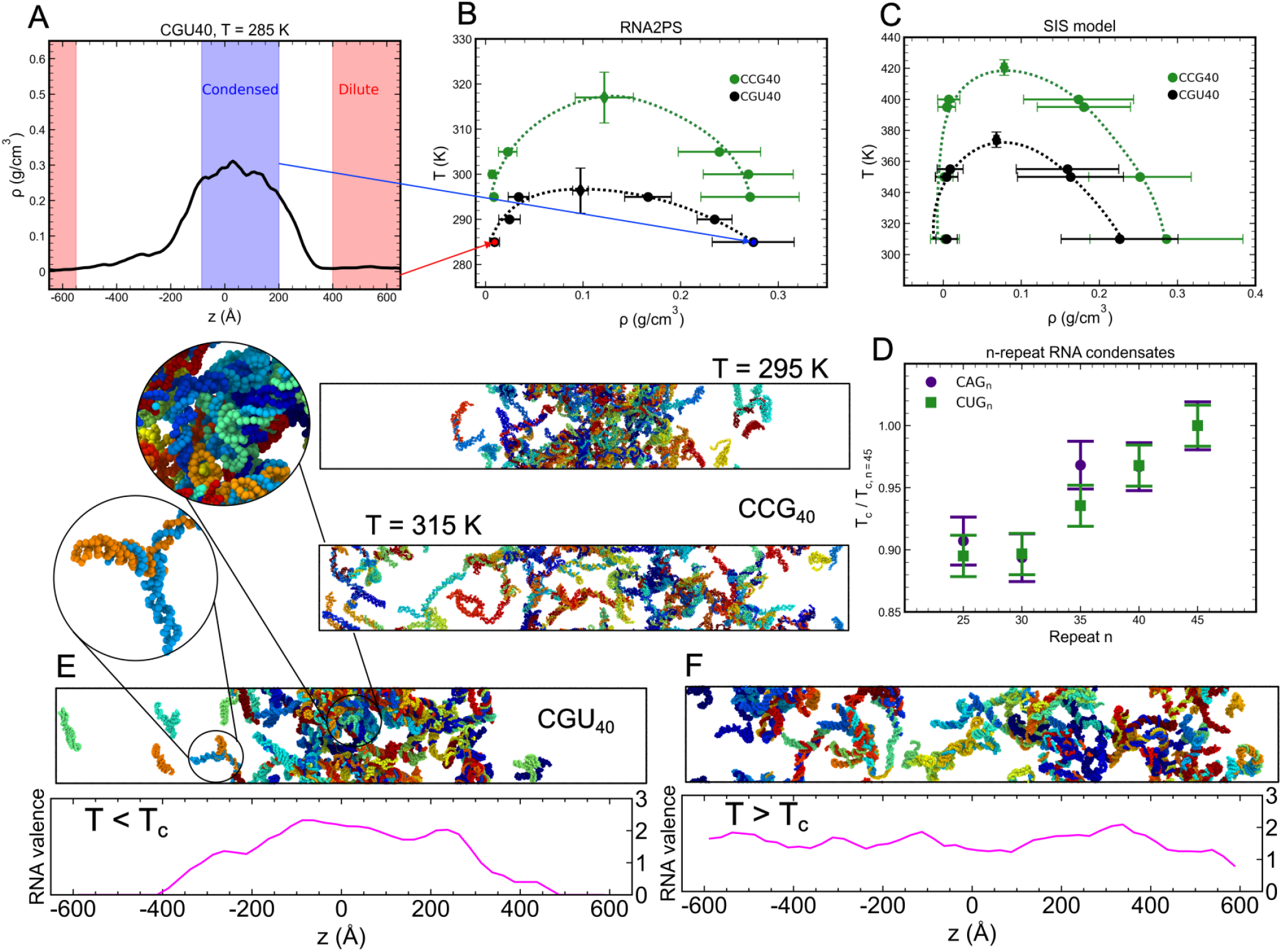
RNA2PS in describing condensate self-assembly. **A**. Density profile of CGU_40_ at T = 285 K in DC simulation. **B**. Phase diagram in the temperature–density plane for the CCG_40_ and CGU_40_ sequences. Critical points are depicted by diamonds. Bottom panels are snapshots of a CCG_40_ simulation below (T = 295 K) and close to (T = 315 K) the critical temperature. **C**. Same phase diagrams as in panel B predicted by the SIS model^63,65^. In both cases, dashed lines are a guide to the eye. **D**. Condensate stability of CAG_*n*_ and CUG_*n*_ repeat sequences increases with repeat length. Normalization constants for the critical solution temperatures of each sequence are T_c, CAG45_ = (259 ± 5) K and T_c, CAG45_ = (302 ± 5) K. The full phase diagrams for these sequences can be found in Fig. S1 in the Supplementary Material. **E**. Simulation snapshots of CGU40 below and **F** above the critical solution temperature. Lower panels represent the valency per RNA molecule in that particular frame. Error bars are calculated from panels **B** and **C** fitting to universal scaling law and law of rectilinear diameters (critical point) and half the difference between maximum and minimum density in condensed or dilute phase; and **D** maximum between simulation grid to identify the critical temperature (5 K) and error from fitting to the universal scaling law.

A key determinant of RNA condensation is chain length, which directly controls the valency and cooperativity of intermolecular interactions^28,29^. To probe this effect, we compute the critical temperatures of CAG_*n*_ and CUG_*n*_ condensates for *n* = 25, 30, 35, 40, and 45 repeats (Fig. 4D). RNA2PS recapitulates this behaviour by reproducing the experimentally observed dependence of condensate stability on *n* repeat length^28,30,63^. As shown in Fig. 4D, the critical temperature is nearly constant for short repeats (*n* ≲ 30), where intermolecular connectivity is limited, and increases with *n* as longer chains enable more extensive multivalent interactions^63^. Remarkably, the normalised critical temperatures of CAG_*n*_ and CUG_*n*_ collapse onto a common trend, suggesting a general scaling behaviour governed primarily by chain length rather than sequence-specific details. Overall, the normalised critical temperature reflects an approximately linear increase with repeat number, highlighting the progressive stabilisation of condensates with increasing the sequence length.

To further characterise the microscopic organisation of RNA within condensates, we analyse the effective valency of RNA molecules, which provides a direct measure of their intermolecular connectivity and multivalent binding capacity. We define molecular valency as the average number of distinct RNA strands with which a given molecule establishes close contacts, computed along the *z*-direction of the simulation box to resolve spatial variations across the interface. Fig. 4E,F shows representative configurations of a CGU_40_ condensate slightly below and above the critical temperature, respectively, together with the corresponding valency profiles (bottom panels). Below *T*_c_, RNA molecules in the dense phase exhibit significantly higher valency (e.g. *>*2) than those at the interface, reflecting the formation of a highly connected network stabilised by intermolecular interactions. This behaviour is consistent with an entropically driven condensation mechanism^92,93^, in which molecules maximise their number of binding partners in the condensed phase. Moreover, it is consistent with previous patchy particle studies on phase separation showing that valency *>*2 is required to establish a percolated liquid network in three dimensions capable to stabilize a condensed phase^94,95^. In contrast, above *T*_c_, the valency becomes nearly uniform across the system (with an average value *<*2 determined by the supersaturation concentration in the solution), indicating the loss of spatial organization and the emergence of a homogeneous phase. These results demonstrate that RNA2PS not only captures RNA condensation at the mesoscopic level, but also recapitulate the underlying changes in molecular connectivity that drive condensate formation.

In Fig. 5, we benchmark the model against experimental cell foci characterisation of RNA condensation, demonstrating its ability to capture sequence-dependent trends across a broad range of repeat sequences with different composition. Fig. 5A shows the fraction of molecules in the largest cluster (FMLC) for non-base-pair-forming RNA repeats at 293 K. RNA2PS exhibits a strong correlation with the experimentally measured fraction of cells forming foci^30^, indicating that the model quantitatively captures intrinsic condensation propensities. Notably, this agreement is achieved by tuning only the base-dependent energy scale of the Wang–Frenkel interaction, with all cross-interactions determined via Lorentz–Berthelot mixing rules and negligible contributions from phosphate beads. In contrast, the SIS model lacks sequence-dependent interactions for non-base-pair-forming sequences, resulting in a nearly flat response that fails to reproduce the experimentally observed variability in phase-separation (Fig. 5B).

**FIG. 5.**
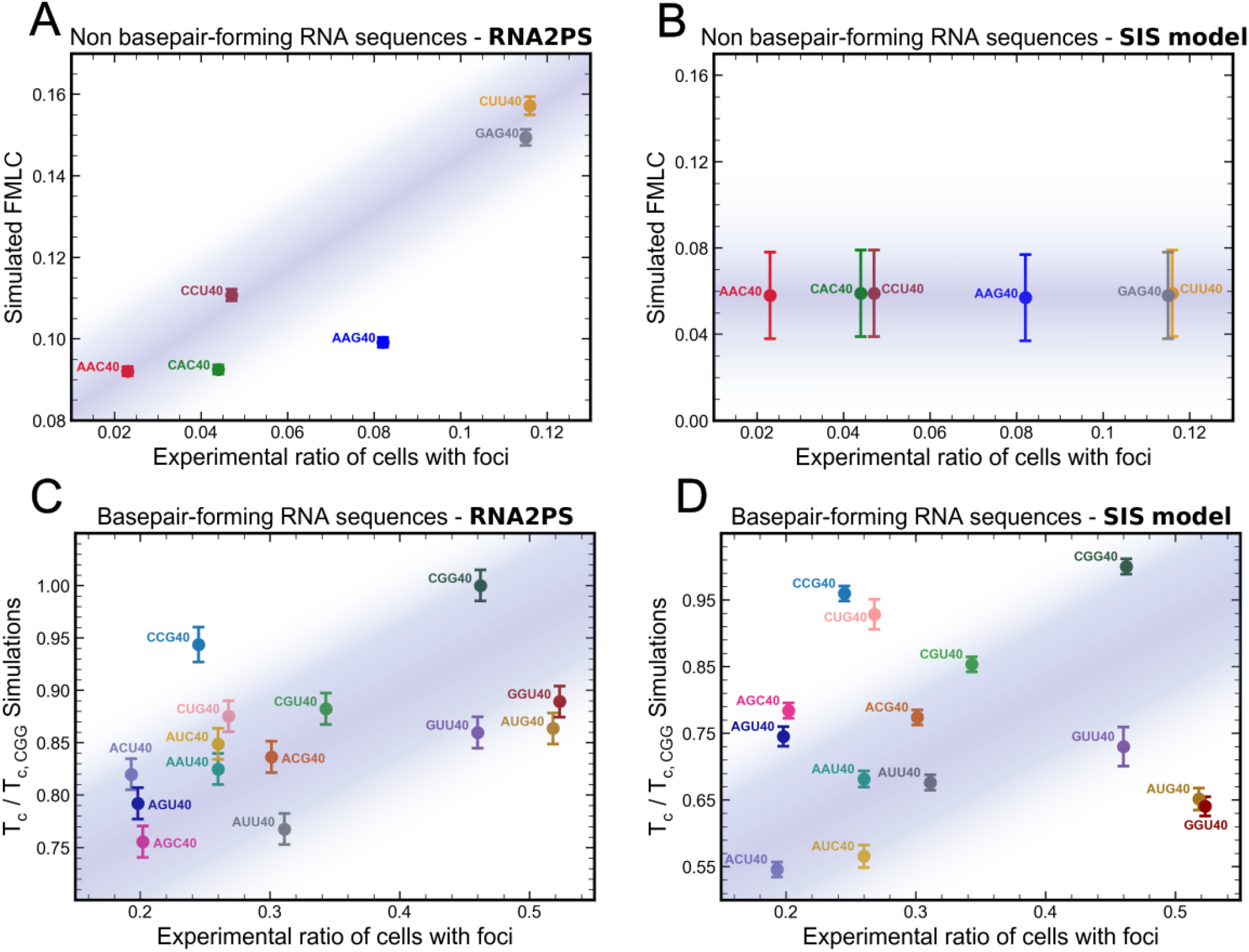
RNA2PS reproduces experimental trends in sequence-dependent RNA condensation. (A) Fraction of Molecules in Largest Cluster (FMLC) of RNA non basepair-forming repeat sequences (at T = 293 K) versus experimental ratio of cells forming foci^30^. (B) Normalized critical temperatures (T_c_) of condensates with basepair-forming RNA repeat sequences correlate with the propensity of cells to express foci. Normalization constant is T_c,CGG_ = (336 ± 5) K. Bottom panels (C) and (D) correspond to the SIS model^63,65^. Normalization constant in (D) is T_c, CGG_ = (438 ± 5) K. Error bars come from (A) and (C) standard error; and (B) and (D) maximum between simulation grid to identify the critical temperature (5 K) and error from fitting to universal scaling law. Phase diagrams for RNA2PS and SIS models can be found in Fig. S2 and Fig. S3 in Supplementary Material, respectively. Shaded regions are a guide to the eye.

We next assess the ability of RNA2PS to predict the stability of condensates formed by base-pair-forming repeats. Fig. 5C shows the correlation between the normalized critical temperature and the experimental propensity to form foci in cells^30^. Notably, RNA2PS reproduces the overall experimental trends as an emergent property, despite base-pairing interactions being calibrated exclusively against melting temperatures and stacking interactions against non-base-pair-forming sequences. The predicted critical solution temperatures for phase-separation span from (255 ± 5) K to (335 ± 5) K. By comparison, the SIS model yields a broader and systematically shifted distribution (240–440 K) and exhibits a significantly weaker correlation with experimental data (Fig. 5D). These results highlight the ability of RNA2PS to link sequence structural properties, thermodynamics and duplex stabilities, as well as condensate phase behaviour within a unified and transferable modelling framework to be combined with protein coarse-grained models^55,61,96**?**^ .

## III. DISCUSSION

In this work, we present RNA2PS, a coarse-grained RNA model with a minimal two-bead-per-nucleotide resolution that enables a unified description of RNA structure, thermodynamics, and phase behaviour. By separating electrostatic interactions, localized on the phosphate bead (although deactivated in this work to study phase-separation under the presence of small divalent cation salt concentration; e.g., 10 mM MgCl_2_^28,29^), from directional base-mediated interactions encoded at the base bead, the model introduces a physically grounded decomposition of RNA interactions. This design preserves nucleotide chemical specificity while providing the flexibility required to capture the competition between intra- and intermolecular contacts that underlies RNA folding and condensation. In contrast to previous two-bead approaches^62^, RNA2PS incorporates explicit sequence-dependent base–base interactions and reproduces the formation of double-helical RNAs, enabling molecular-level insight into both isolated RNA structures and collective multi-chain pure RNA condensation.

A key feature of RNA2PS is its ability to reconcile the formation of stable double helices with the conformational flexibility of single disordered strands. In RNA2PS, the geometry of base pairing is encoded through a multi-body interaction, ensuring that double-helical conformations emerge naturally while preventing unphysical overcoordination and simultaneous binding to multiple partners^63^. At the same time, the spatial separation between backbone and base beads introduces intrinsic directionality and excluded-volume constraints, which restrict the relative orientation and proximity of interacting bases. This effectively captures the configurational and entropic cost associated with base pairing and stacking, and limits the number of favourable contacts that each nucleotide can establish. This balance is essential: intramolecular base pairing stabilizes folded conformations, whereas intermolecular interactions promote multivalency and biomolecular condensate formation.

Such interplay is directly reflected in the predicted structural topology of RNA double-strands (Fig. 2) and the thermodynamic behaviour of the RNA2PS model. RNA2PS quantitatively reproduces the melting temperatures of short RNA sequences in dilute conditions (Fig. 3), with a standard deviation of 17.5 K from experimental values, while simultaneously predicting critical temperatures for RNA condensates within biologically relevant ranges (260-335 K; Figs. 4 and 5). Achieving this level of consistency across such distinct properties highlights the model’s ability to capture the underlying free-energy landscape governing RNA phase behaviour. In RNA2PS, condensation of RNA solutions in presence of implicit cation divalent ions emerges from a subtle balance between enthalpic gain associated with base pairing and stacking interactions as well as from entropic contributions arising from multivalent intermolecular connectivity. The formation of interchain contacts increases the number of accessible binding configurations at the network level, effectively enhancing the degeneracy of the condensed state. The two-bead architecture plays a central role in regulating this balance: by geometrically constraining base pairing and limiting the number of simultaneous interactions per nucleotide, it prevents unphysical overcoordination and enforces a finite valency. As a result, the model avoids the overstabilization of the condensed phase predicted by other models^63^ and preserves a more realistic competition between intra- and intermolecular interactions. This is particularly relevant for repeat-expansion RNAs, where small variations in sequence composition and length lead to measurable differences in condensation propensity^30^. Importantly, this level of consistency is achieved within a minimal representation, avoiding the need for higher-resolution models^74^, which, although accurate, are often limited by computational cost to large scale systems.

RNA2PS further captures sequence-dependent trends in condensation propensity for both base-pair-forming and non-base-pair-forming RNA repeat sequences (Fig. 5). For base-pair-forming sequences, the predicted critical temperatures follow the expected stability trends^65,92^, reflecting the interplay between interaction strength, sequence-dependent base pairing energetics, and the number of accessible binding configurations. This framework explains the enhanced stability of sequences such as CCG_40_ and CGG_40_, which maximize both base-pairing propensity and multivalency. Moreover, the model highlights the role of non-canonical and dispersive interactions, particularly in sequences containing G–U motifs^29^, in modulating condensate stability beyond canonical Watson–Crick pairing. Deviations observed for specific sequences in cell foci may indicate additional contributions not captured within an RNA-only description, such as protein-mediated interactions in highly multicomponent cellular environments^30^.

The level of coarse-graining and dynamical properties of RNA2PS are well matched to those of state-of-the-art sequence-dependent coarse-grained protein models commonly used to study biomolecular condensates, such as the Mpipi, CALVADOS, and HPS families^55,60,61,96,97^. This compatibility will enable the direct simulation of RNA–protein mixtures with explicit RNA base-pairing. In this context, the RNA2PS model, once its cross-interactions with protein force fields are optimized—a task beyond the scope of this work—can provide a robust framework to explore RNA-driven regulation of phase separation^11,22^, including processes such as stress granule self-assembly^98^ or RNA-mediated modulation of protein condensation^20^ among others^25,99^. RNA–protein assemblies are recognized as central to intracellular organization and regulation, and their dysregulation is increasingly linked to multiple diseases^29,100^, including certain cancers, where aberrant condensate formation can alter transcriptional programs and cellular identity^101,102^. In this context, tumor-associated condensates provide a particularly relevant example in which changes in RNA– or DNA–protein interactions may directly impact phase behaviour and function. A physically consistent multi-component model can be therefore essential to dissect the molecular principles governing condensate formation and its dysregulation in pathological states.

## IV. MATERIALS AND METHODS

### A. Model force field

We present the RNA2PS, a coarse-grained computational model with two-beads-per-nucleotide resolution to simulate RNA self-condensation. One bead is located at the geometric centre of mass of the phosphate+ribose groups (henceforth phosphate bead, P), and the other at 4 Å in the direction of the vector connecting P to the centre of the nitrogenous base (henceforth base bead, B). The total potential energy of the system is defined as the sum of bonded and non-bonded contributions.

Bead connectivity is enforced through harmonic bond potentials of the form

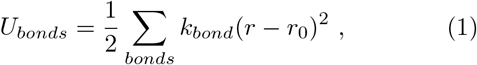

being *r* the bond distance and *r*_0_ its equilibrium value. Bonds are established between consecutive phosphate and base beads, as well as between bases and phosphates linked by a phosphodiester bond. The rigidity is modelled by means of an angular potential with the form

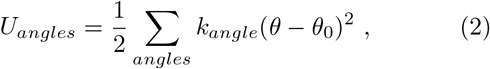

where *θ* is the angle formed by any three consecutive base beads and *θ*_0_ its equilibrium value. Table I contains all intramolecular interaction parameters used in RNA2PS. Base-pairing interactions are described by means of a multibody potential^63^ with the form

**TABLE I.** Values of the intramolecular bonded interaction parameters. P and B denotes backbone and nucleotide beads, respectively.

| Parameter | Value |
| --- | --- |
| $r_0(\text{P-P}) / \text{\AA}$ | 5.2 |
| $r_0(\text{B-B}) / \text{\AA}$ | 3.8 |
| $r_0(\text{P-B}) / \text{\AA}$ | 4.0 |
| $k_{bond}(\text{P-P}) / \text{kcal mol}^{-1} \text{\AA}^2$ | 8.8 |
| $k_{bond}(\text{B-B}) / \text{kcal mol}^{-1} \text{\AA}^2$ | 1.5 |
| $k_{bond}(\text{P-B}) / \text{kcal mol}^{-1} \text{\AA}^2$ | 6.95 |
| $\theta_0 / \text{degrees}$ | 150 |
| $k_{angle} / \text{kcal mol}^{-1}$ | 6.05 |

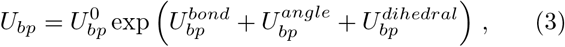

where 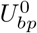 sets the overall strength of the base-pairing interaction and depends on sequence context, and

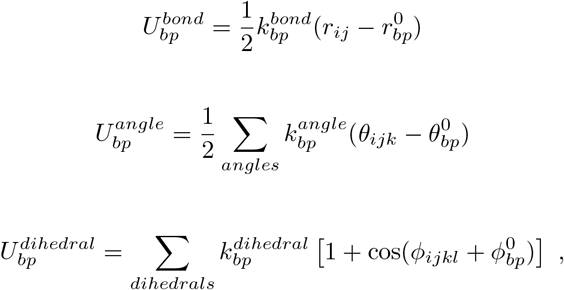

where 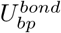 depends on the distance between the beads forming the base pairing (*i* and *j*); 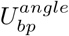 includes contributions from four angles centred at *i* and *j*, each involving the opposing base bead and one of its neighbouring phosphate beads; and 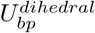 accounts for the two dihedrals angles defined by *i, j*, and the adjacent base beads along the strand. In addition, 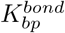 depends on the specific base identity. Tables II and III summarise all model parameters in Eq. (3). The potential is only applied to bead pairs *i* and *j* that (i) have compatible biochemical identities, (ii) are separated by least six nucleotides along the same strand, (iii) lie within a cut-off distance of 8 Å, and (iv) are not located at the termini of the molecule.

**TABLE II.** Values of the intermolecular bonds corresponding to the base pairing parameters.

| Parameters | Pair Type |  |  |
| --- | --- | --- | --- |
|  | AU | CG | GU |
| $r_{bp}^0$ / Å | | 6.9 | |
| $k_{bp}^{bond}$ / kcal mol<sup>-1</sup> Å<sup>2</sup> | 1.9 | 1.15 | 1.80 |

**TABLE III.**
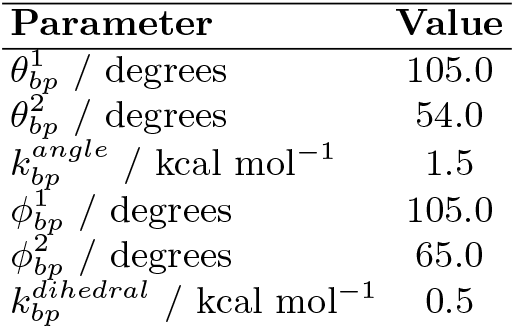
Values of the intermolecular angles and dihedrals corresponding to the base pairing parameters.

Cross-stacking interactions are represented by a Wang-Frenkel potential^75^ of the form

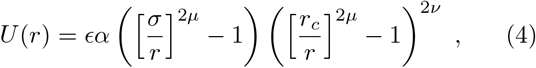

where

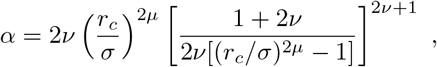

and

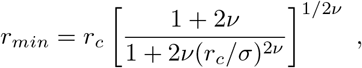

where *r*_*c*_ is the cut-off radius, *ν* = 1, *µ* = 3 and the energy (*ϵ*) and distance (*σ*) parameters are given in Table IV. Cross interactions follow Lorentz-Berthelot mixing rules.

**TABLE IV.** Wang-Frenkel parameters. R_*c*_ is selected for 3*σ*_*i*_.

| Parameters | Bead Type |  |  |  |  |
| --- | --- | --- | --- | --- | --- |
|  | P | A | U | C | G |
| $\epsilon$ / kcal mol<sup>-1</sup> | 6.954 | 6.430 | 5.958 | 6.531 | 6.033 |
| $\sigma$ / Å | 0.0100 | 0.1998 | 0.5827 | 0.2498 | 0.5827 |

Finally, electrostatic interactions between phosphate beads are modelled using a Yukawa potential. In this work, all properties are evaluated in the high-salt regime, where the Debye screening length is effectively negligible and electrostatic contributions are strongly screened. Nevertheless, including this term is essential to capture the interplay between RNA and proteins in biomolecular condensates.

### B. Simulation of RNA crystal structures

We carried out molecular dynamics simulations to determine the optimal equilibrium parameters for bond, angle and dihedral potentials describing both base-pairing interactions and intramolecular RNA structure. Starting from experimental PDB structures, atoms were mapped onto the RNA2PS representation by placing the backbone bead at the centre of mass of the ribose+phosphate and the base bead along the vector connecting the backbone bead to the centre of mass of the nitrogenous base, at a distance of 4Å. For each sequence, independent simulations were performed at T=250*K* to mimic the stability of the crystal structure at low temperature. Each simulation run for 1 *µ*s and included 64 replicas of the structure under dilute conditions (∼10.5 mM) to prevent intermolecular interactions. Bond lengths, angles and dihedrals were extracted from both the mapped PDB structures and the simulation trajectories, and their distributions were compared. This procedure was iteratively refined using a basin-hopping algorithm to minimise the error. In total, a dataset of 25 RNA PDB structures was used.

Molecular dynamics simulations were performed to estimate the melting temperature (T_m_) or RNA duplexes. For each sequence, independent simulations were carried out at temperatures ranging from 280 K to 355 K, using 15 K intervals. Each simulation was run for 1 *µ*s, and configurations were saved every 0.1 ns for subsequent analysis. The systems contained 128 RNA strands (64 duplexes) placed in a cubic box of dimensions 200×200×200 Å^3^, corresponding to RNA strand concentration of 26.6 mM. From the resulting trajectories, the number of dissociated strands was computed as a function of time and temperature. The fraction of dissociated strands was used to construct thermal denaturation curves, and the melting temperatures were defined as the temperature at which 50% of the strands are dissociated. This value was obtained by fitting the data to a sigmoidal function of the form 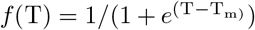, where T_m_ is the melting temperature. This procedure was applied to a total of 33 RNA sequences, including 5 sequences containing mismatches.

### C. Simulation of RNA condensates

We simulate both base-pairing and non-base-pairing condensates using the direct coexistence method^89,95^. Condenstates composed of 100 molecules were prepared in an orthorhombic box with dimensions 210.50×210.50×1376.30 Å^3^, corresponding to an overall density of approximately 0.07 *g/cm*^3^. The initial configuration was constructed at higher density of 0.40 *g/cm*^3^ to promote condensation. For non-base-pairing systems, simulations were carried out at 293 *K* over 2 *µs*, and the fraction of molecules in the largest cluster was averaged over the final 1.6 *µs*. For base-pairing systems, configurations were first heated to 350 K for 1 *ns* to disrupt base pairing and avoid kinetic trapping. Simulations at the target temperature were then performed for 3 *µs*, with only the last 1.2 *µs* used for analysis. In all cases, the integration time step was set to 10 *fs*, and configuration were saved every 2 *ns*.

### D. Physical observables

We characterise the propensity of non-base-pairing condensates by measuring the fraction of molecules in the largest cluster at T = 293 *K*. Clusters are defined as sets of molecules in which each molecule lies within 8 Å of at least one other molecule in the cluster. The distance between two molecules is defined as the minimum distance between any pair of beads belonging to each molecule.

The stability of base-pairing condensates is assessed by analysing their critical temperatures. We determine the critical point (*ρ*_*c*_, *T*_*c*_) of each system using the universal scaling relations near the critical point^91^

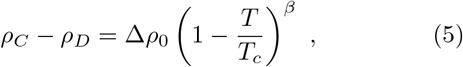

where *ρ*_*C*_ and *ρ*_*D*_ are the densities of the condensed and dilute phase, respectively, *β* ≈ 0.325 is the critical exponent, and Δ*ρ*_0_ is a fitting parameter. In addition, the law of rectilinear diameters^91^ is used,

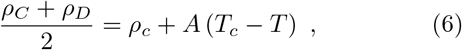

being *A* is a fitting parameter. The densities *ρ*_*C*_ and *ρ*_*D*_ are obtained by averaging appropriate regions of the density profiles at each temperature. All fits are performed using three data points below the critical temperature, with temperature spacing of 5-10 *K* between consecutive points.

## V. DATA AVAILABILITY

LAMMPS implementation files have been made available on Github https://github.com/pllombar/RNA2PS.

## VI. ACKNOWLEDGEMENTS

A. R. T. acknowledges funding from the MICIU under the Juan de la Cierva fellowship (JDC2024-053759-I). P.L. and J.L-M. acknowledges funding from the European Union’s Horizon 2020 research and innovation program (grant agreement 101160499 to J. R. E). J.R. acknowledges funding from the Spanish ministry of Science and Innovation (project number PID2022-136919NB-C32). R.C.-G. acknowledges funding the UK Research Innovation (UKRI) Engineering and Physical Sciences Research Council (EPSRC) [EP/Z002028/1], following funding from the European Research Council (ERC) Consolidator Grant “ChromatinDroplets” under the European Union’s Horizon Europe research and innovation programme. J. R. E. acknowledges funding from Emmanuel College, the University of Cambridge, the Ramon y Cajal fellowship (RYC2021-030937-I), the Spanish scientific plan and committee for research reference PID2022-136919NA-C33, and the ERC under the European Union’s Horizon Europe research and innovation program (grant agreement no. 101160499).

This work has been performed using resources provided by the Cambridge Tier-2 system operated by the University of Cambridge Research Computing Service (http://www.hpc.cam.ac.uk) funded by EPSRC Tier-2 capital grant EP/P020259/1-CS170. This work has also been performed using resources provided by Archer2 (https://www.archer2.ac.uk/) funded by EPSRC Tier-2 capital grant EP/P020259/e829. The authors also thankfully acknowledge RES computational resources provided by Mare Nostrum 5 through the activity FI-2025-3-0065.

## Notes

### Competing Interest Statement

The authors have declared no competing interest.

