## Supporting Information for "RNA2PS: A sequence-specific coarse-grained RNA model linking structure, thermodynamics and phase separation"

Juan Luengo-Márquez[†]

*Department of Physical Chemistry, Universidad Complutense de Madrid,*
*Av. Complutense s/n, Madrid 28040, Spain*

Jorge Ramirez

*Department of Chemical Engineering,*
*Universidad Politécnica de Madrid,*
*José Gutiérrez Abascal 2, 28006, Madrid, Spain.*

Alberto Ocana

*Experimental Therapeutics Unit, Hospital Clínico San Carlos (HCSC),*
*Instituto de Investigación Sanitaria San Carlos (IdISSC), Madrid, Spain and*
*PhAsIca Biosciences S.L, Calle Velázquez, 27, 28001 Madrid, Spain*

Rosana Collepardo-Guevara

*Yusuf Hamied Department of Chemistry,*
*University of Cambridge, Lensfield Road,*
*Cambridge CB2 1EW, United Kingdom and*
*Department of Genetics, University of Cambridge, Cambridge, CB2 3EH*

Jorge R. Espinosa [a]

*Department of Physical Chemistry, Universidad Complutense de Madrid,*
*Av. Complutense s/n, Madrid 28040, Spain*

*Instituto Pluridisciplinar, Universidad Complutense de Madrid,*
*P.º de Juan XXIII, 1, Moncloa - Aravaca, 28040 Madrid, Spain*
*Yusuf Hamied Department of Chemistry, University of Cambridge,*
*Lensfield Road, Cambridge CB2 1EW, UK and*
*PhAsIca Biosciences S.L, Calle Velázquez, 27, 28001 Madrid, Spain*
(Dated: April 24, 2026)

[a]

[b]

†These authors contributed equally

| **Base** | **Bead type** | (Nearest-Neighbour sequence) | |
|---|---|---|---|
| A | A1 (XAX) | A5 (5AX) | |
| | A2 (AAX) | A6 (5AA) | |
| | A3 (XAA) | A7 (3AX) | |
| | A4 (AAA) | A8 (3AA) | |
| C | C1 (XCX) | C5 (5CX) | |
| | C2 (CCX) | C6 (5CC) | |
| | C3 (XCC) | C7 (3CX) | |
| | C4 (CCC) | C8 (3CC) | |
| G | G1 (XGA) | G6 (AGG) | G11 (5GG) |
| | G2 (XGX) | G7 (GGA) | G12 (3GX) |
| | G3 (AGX) | G8 (AGA) | G13 (3GG) |
| | G4 (GGX) | G9 (GGG) | G14 (5GA) |
| | G5 (XGG) | G10 (5GX) | G15 (3GA) |
| U | U1 (XUC) | U6 (CUU) | U11 (5UU) |
| | U2 (XUX) | U7 (UUC) | U12 (3UX) |
| | U3 (CUX) | U8 (CUC) | U13 (3UU) |
| | U4 (UUX) | U9 (UUU) | U14 (5UC) |
| | U5 (XUU) | U10 (5UX) | U15 (3UC) |

**TABLE S1:** Bead types and nearest-neighbour sequences grouped by base.

| PDB code | Reference |
|---|---|
| 1A4D | https://doi.org/10.1016/S0969-2126(97)00311-0 |
| 1DQF | http://doi.org/10.1017/S135583820000090X |
| 1KKA | https://doi.org/10.1016/S0022-2836(02)00382-0 |
| 1KXK | http://doi.org/10.1126/science.1069268 |
| 1RNA | http://doi.org/10.1016/0022-2836(89)90010-7 |
| 259D | http://doi.org/10.1021/bi9607214 |
| 2A43 | http://doi.org/10.1021/bi051061i |
| 2KX8 | https://doi.org/10.1016/j.jmb.2010.08.042 |
| 2L2J | https://doi.org/10.1016/j.cell.2010.09.026 |
| 3MEI | http://doi.org/10.1107/S0907444910050900 |
| 4E59 | https://doi.org/10.1093/nar/gks557 |
| 4JAB | http://doi.org/10.2210/pdb4jab/pdb |
| 4KYY | https://doi.org/10.2210/pdb4KYY/pdb |
| 5BTM | https://doi.org/10.1021/acs.biochem.5b00551 |
| 6XWJ | https://doi.org/10.1093/nar/gkaa465 |
| 7WIB | http://doi.org/10.1093/nar/gkac1257 |
| 8AMJ | http://doi.org/10.1261/rna.079414.122 |
| 8K2Z | https://doi.org/10.1038/s41421-024-00761-1 |
| 8K8B | https://doi.org/10.1016/j.bbrc.2023.149327 |
| 8ZAU | https://doi.org/10.1093/nar/gkae1218 |
| 9BXE | https://doi.org/10.1016/j.bpj.2025.11.026 |
| 9CSO | https://doi.org/10.1073/pnas.2505720122 |
| 9CSQ | https://doi.org/10.1073/pnas.2505720122 |
| 9DE6 | https://doi.org/10.1038/s41467-025-57519-w |
| 9I8B | https://doi.org/10.1093/nar/gkaf550 |

**TABLE S2:** PDB entries and their corresponding references used for structural optimization of RNA2PS

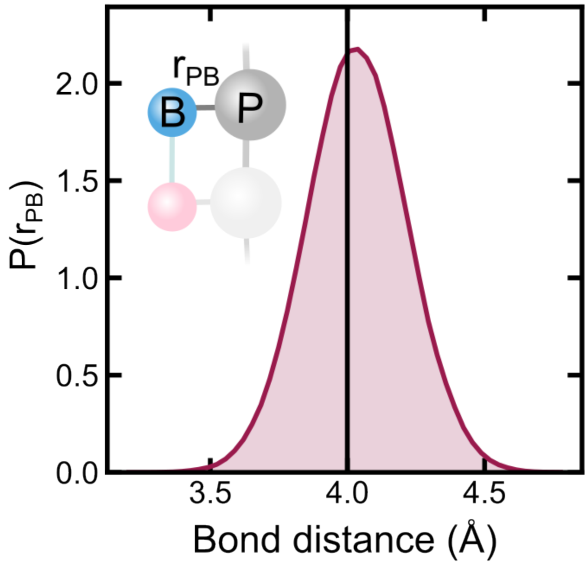


**FIG. S1:** Distribution of intramolecular P-B bond distance. Black line indicates the distance fixed in the mapping grom the PDB crystal structures.

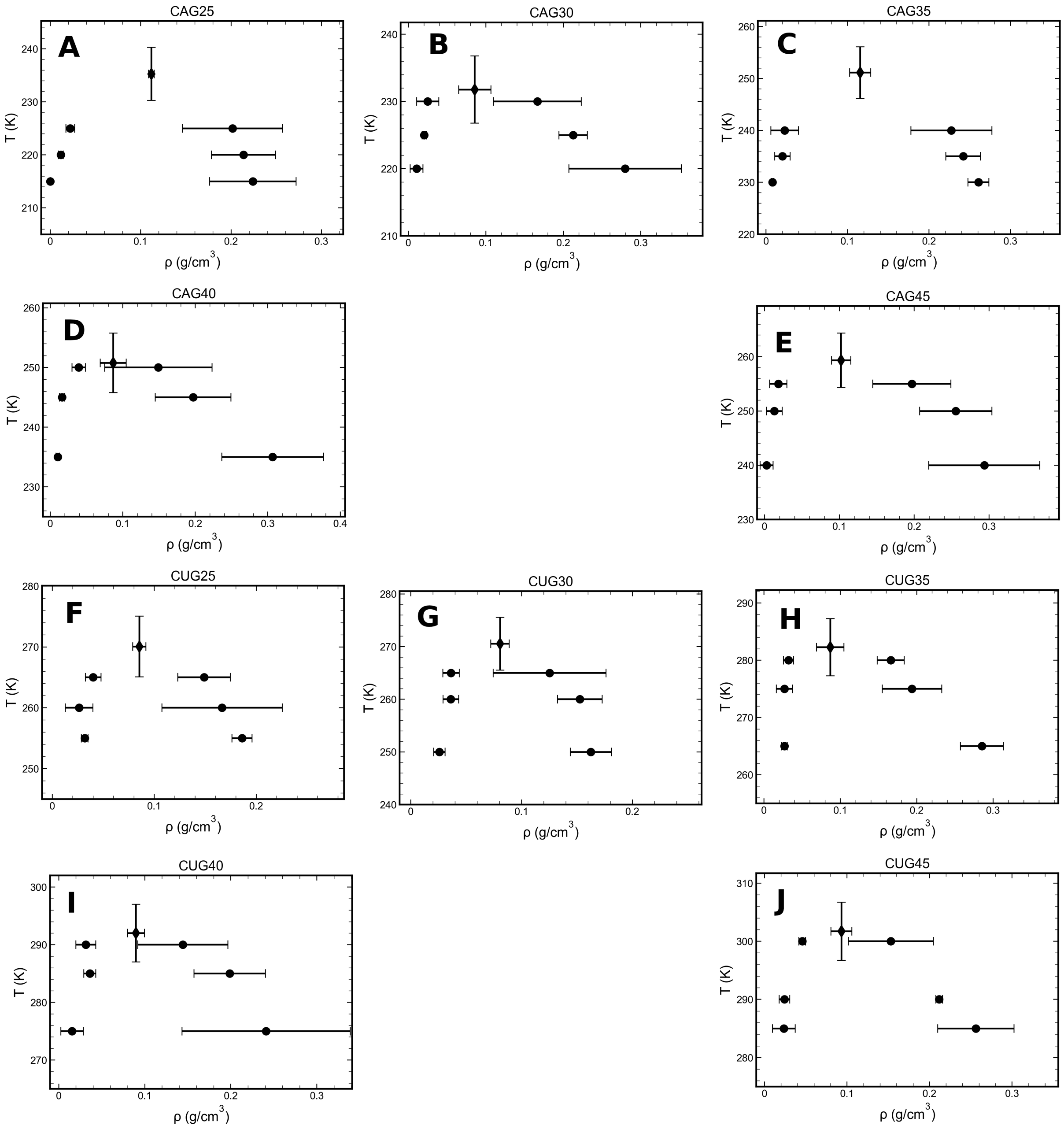


**FIG. S2:** Phase diagrams of RNA2PS RNA repeat condensates.

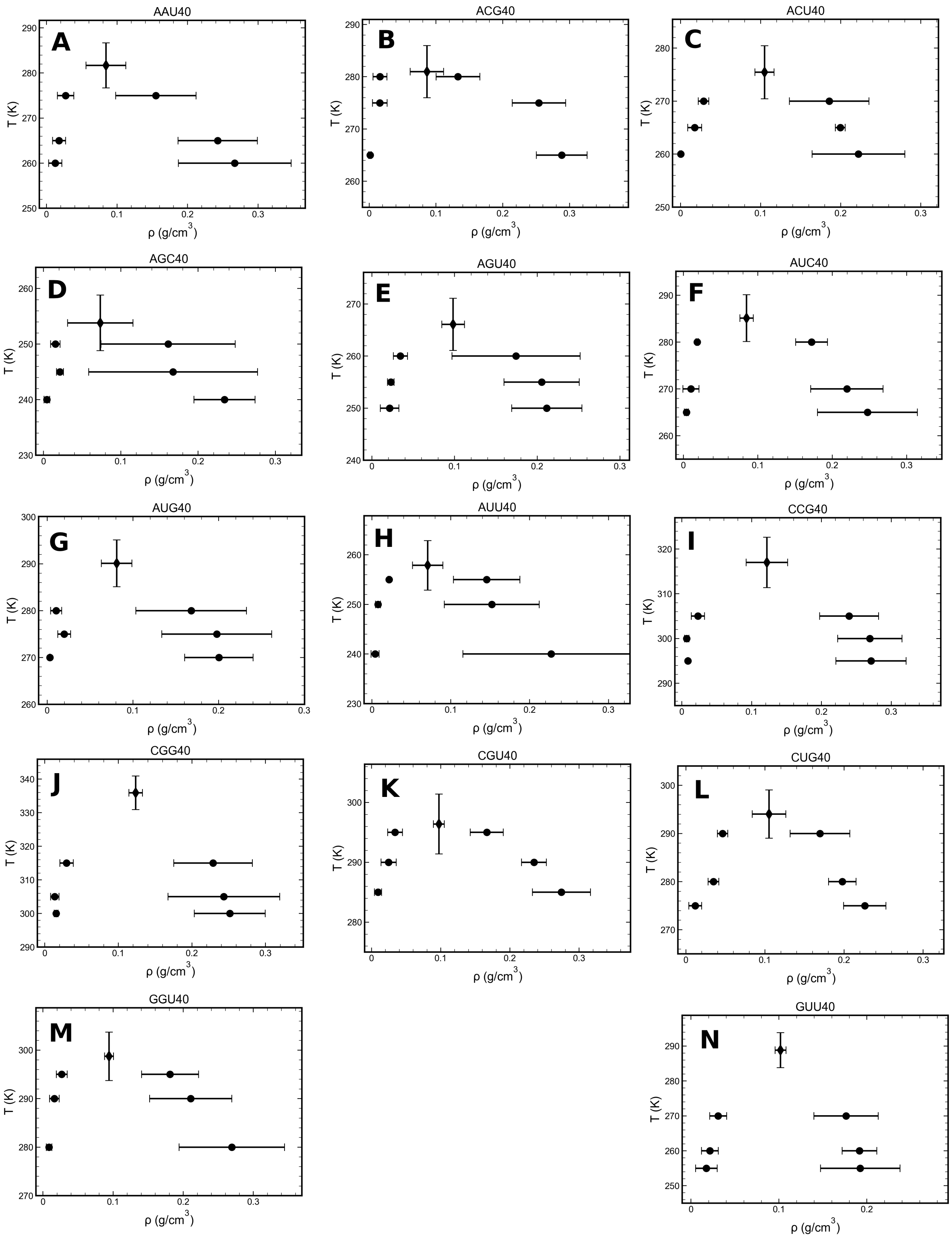


**FIG. S3:** Phase diagrams of RNA2PS condensates.

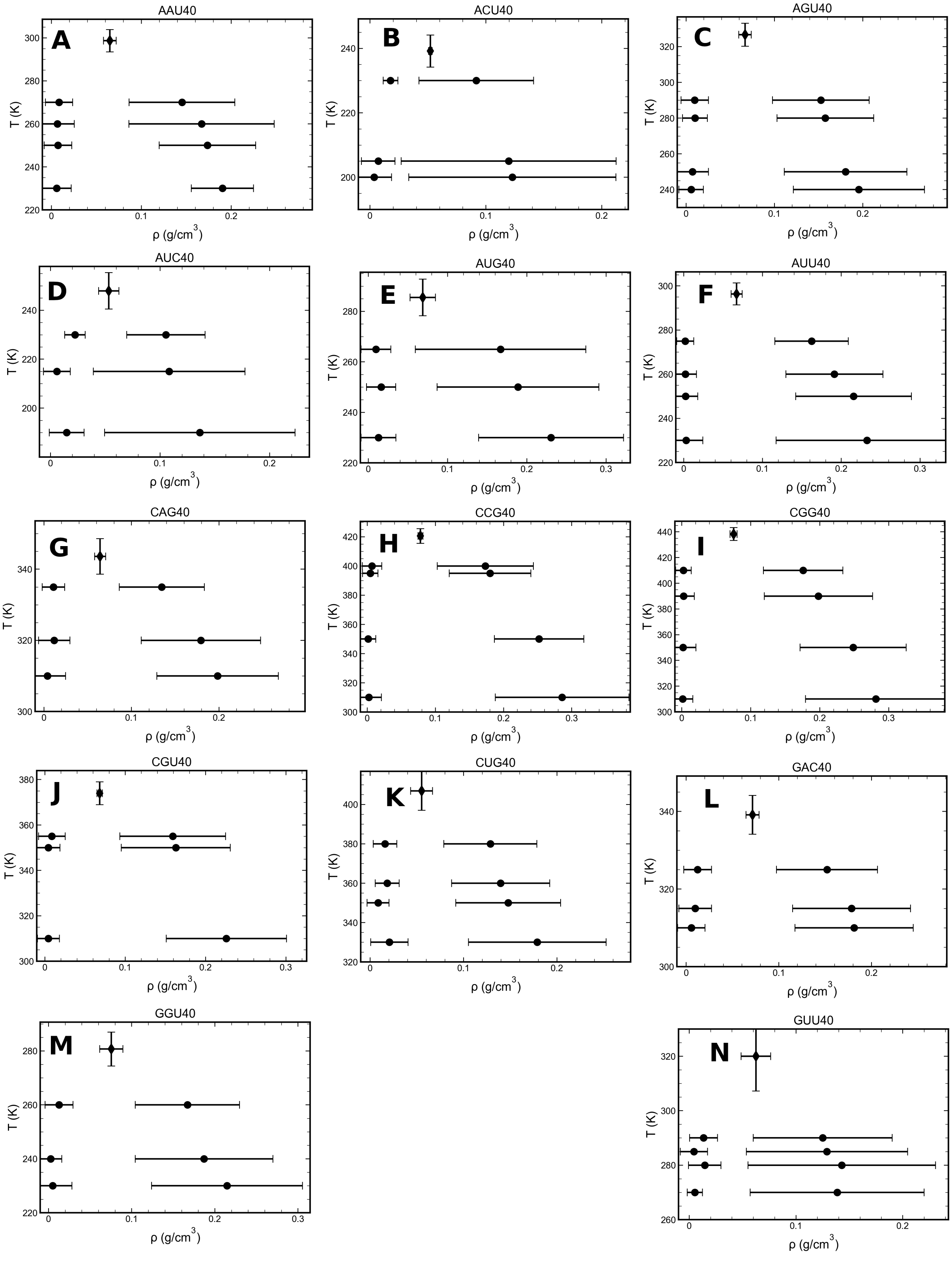


**FIG. S4:** Phase diagrams of SIS condensates.
